# Convergent remodeling of the gut microbiome is associated with host energetic condition over long-distance migration

**DOI:** 10.1101/2022.11.30.518533

**Authors:** Brian K. Trevelline, Daniel Sprockett, William V. DeLuca, Catherine R. Andreadis, Andrew H. Moeller, Christopher Tonra

## Abstract

The gut microbiome can be thought of as a ‘forgotten organ’, owing to its profound effects on host phenotypes. Long-distance migratory birds are capable of adaptively modulating their physiology, raising the hypothesis that the microbiome of migratory birds may undergo a parallel remodeling process that helps to meet the energetic demands of long-distance migration. To test this hypothesis, we investigated changes in gut microbiome composition and function over the fall migration of a Neotropical-Nearctic migratory Blackpoll Warbler (*Setophaga striata*), which exhibits one of the longest known autumnal migratory routes of any songbird and rapidly undergoes extensive physiological remodeling during migration. Overall, our results showed that the Blackpoll warbler microbiome differed significantly across phases of fall migration. This pattern was driven by a dramatic increase in the relative abundance of Proteobacteria, and more specifically a single ASV belonging to the family Enterobacteriaceae. Further, blackpolls exhibited a progressive reduction in microbiome phylogenetic diversity and within-group variances over migration, indicating convergence of microbiome composition among individuals during long-distance migration. Metagenomic analysis revealed that the gut microbiome of staging blackpolls was enriched in bacterial pathways involved in vitamin, amino acid, and fatty acid biosynthesis, as well as carbohydrate metabolism, and that these pathways were in turn positively associated with host body mass and subcutaneous fat deposits. Together, these results provide evidence that the gut microbiome of migratory birds may undergo adaptive remodeling to meet the physiological and energetic demands of long-distance migration.

## INTRODUCTION

The microbial communities of the intestinal tract, known as the gut microbiome, can profoundly influence animal phenotypes through a myriad of effects on host physiology (McFall-Ngai et al., 2013). The gut microbiome is shaped by a variety of extrinsic environmental and intrinsic host factors (Ley et al., 2008a; Muegge et al., 2011; Rothschild et al., 2018a), which can influence host phenotypes through their effects on microbiome functions (Cho & Blaser, 2012; Clemente et al., 2012). However, the vast majority of research on the effect of the gut microbiome on host phenotypes has been derived from humans and model organisms under highly static conditions (Colston & Jackson, 2016; Pascoe et al., 2017). We now recognize that there is immense variation in host microbiota both within and between individuals, across habitats, and through time, and that these differences may reflect differences in host adaptive capacity and fitness in nature (Alberdi et al., 2016; Suzuki, 2017). Therefore, understanding the relationship between the gut microbiome and host phenotypes through space and time is essential to our understanding of wild vertebrate ecology and evolution (Moran et al., 2019; Trevelline, Fontaine, et al., 2019).

Animals faced with extreme energetic or physiological challenges are ideal natural models for investigating potentially adaptive microbiome-mediated host phenotypes. The gut microbiome of wild vertebrates can confer the ability for their hosts to tolerate seasonal temperature extremes (Fontaine et al., 2022), consume diets which would otherwise be toxic (Kohl et al., 2014, 2016), resist parasitic infections (Knutie et al., 2017), and balance nitrogen during fasting (Wiebler et al., 2018). Further, seasonal differences in host diet can trigger the remodeling of gut microbiome in a way that influences host phenotypes such as intestinal epithelial structure, nitrogen balance, immune function, and metabolism (Carey & Assadi-Porter, 2017). Many of these microbiome-mediated host phenotypes could influence vertebrate fitness in the wild, and thus adaptive microbiome functions may be favored by natural selection (Fontaine & Kohl, 2020; Groussin et al., 2017; Henry et al., 2021; Ley et al., 2008b; Lynch & Hsiao, 2019; Zilber-Rosenberg & Rosenberg, 2008).

Neotropical-Nearctic migratory birds annually complete some of the most energetically-demanding long-distance movements in the animal kingdom and can adaptively modulate aspects of their physiology to meet the extreme energetic demands of migration (McWilliams & Karasov, 2001). At the conclusion of a months-long stationary breeding period, migrants begin depositing protein (in the form of muscle and organ tissue) and fat, which provide the primary source of metabolic energy for long-distance migration (Jenni & Jenni-Eiermann, 1998). In general, migration is initially characterized by the depletion of these energy reserves during short overnight flights followed by daytime ‘refueling’ of subcutaneous fat stores and recovery of muscle and organ tissue at intermediate “stopover” locations (McWilliams & Karasov, 2001; Schwilch et al., 2002a). Nearctic-neotropical migrants repeat this depletion-recovery cycle of energy reserves at stopover sites until they reach a natural barrier (*e*.*g*., ocean, desert) where they are faced with the extreme challenge of a multi-day, non-stop flight over thousands of kilometers without food, water, or rest. To overcome this energetic challenge, some migrants undergo a weeks-long form of stopover known as ‘staging’ – a period characterized by extreme hyperphagia, increased muscle and digestive system mass, and a rapid accumulation of subcutaneous fat stores in the weeks prior to long bouts of flight (Warnock, 2010). The ability to adaptively modulate phenotypes that enhance migratory performance is a unifying characteristic among migratory birds (McWilliams & Karasov, 2001; Piersma, 1998), and therefore is considered a key adaptation in the evolution of the migration (Alerstam et al., 2003).

The gut microbiome has been referred to as the ‘forgotten organ’, owing to its profound effects on host physiology (O’Hara & Shanahan, 2006). This raises the possibility that, like other organs, the microbiome of migratory birds undergoes a potentially adaptive remodeling that helps to meet the energetic demands of long-distance migration. While limited to just a few descriptive studies, previous work has demonstrated that gut microbiome community composition of migratory birds differs between breeding and wintering habitats (Skeen et al., 2021; Wu et al., 2018), and is responsive to the timing and duration of migratory stopover (Lewis et al., 2016, 2017; Thie et al., 2022). Further, these studies have demonstrated that migration can lead to the enrichment of certain microbial taxa (e.g., Corynebacteria and Proteobacteria; Risely et al., 2017, 2018; Skeen et al., 2021; Turjeman et al., 2020). Importantly, the functional capacity of the microbiome is directly related to its composition (Burke et al., 2011), and thus these findings suggest that the demands of migration exert strong selective pressures that could potentially lead to an adaptive enrichment of microbiome functions that help the host overcome the extreme energetic demands of long-distance migration. However, whether variation in microbiome composition over space and time is associated with the enrichment of microbial functions that could enhance host energy metabolism remains unknown.

In this study, we investigated changes in gut microbiome composition and function over the fall migration of a Neotropical-Nearctic migratory passerine, the Blackpoll Warbler (*Setophaga striata*; hereafter blackpolls). Each fall, blackpolls complete one of the longest known migratory routes of any songbird (∼8,000 km), from breeding sites in Western Canada to wintering habitats in northern South America (Fig. S1; DeLuca et al., 2015, 2020). Similar to other species of migratory songbirds, blackpoll migration begins following a months-long stationary breeding phase and is initially characterized by short overnight flights to inland stopover sites where they recuperate their energy reserves (DeLuca et al., 2020; Morris et al., 2016). This process is repeated over a period of several months as blackpolls progress southeast across North America to the Atlantic Ocean, where they are faced with a non-stop transoceanic migration (up to ∼3,500 km/∼80 hours) to wintering sites in South America (DeLuca et al., 2015, 2020; Williams et al., 1978). To successfully complete this energetically-demanding feat, blackpolls along the Atlantic coast undergo a weeks-long process known as pre-migration staging characterized by an extensive physiological transformation that includes a rapid increase in body mass and subcutaneous fat deposits (DeLuca et al., 2020), which are important metrics of energetic condition in blackpolls (Smetzer et al., 2017). To test the hypothesis that the gut microbiome of can be adaptively remodeled to meet the demands of migration, we characterized changes in gut microbiome composition and function from fecal samples collected at multiple sites within each distinct phase of the Blackpoll warbler’s North American fall migratory route. We predicted that the gut microbiome of staging blackpolls would be enriched in microbial pathways that could enhance blackpoll energy metabolism during long-distance migration, and that these pathways would be associated with metrics of their energetic condition.

## MATERIALS AND METHODS

### Sample and data collection

In collaboration with several research groups and avian monitoring stations across North America, blackpoll fecal samples were collected for microbiome analysis at multiple locations within three distinct phases of their fall migration: breeding, migratory stopover, and pre-migration staging (Figure S1; Supplemental Data 1). Fecal samples collected from breeding blackpolls in Western Canada were categorized as “breeding” (Yukon and British Columbia). Blackpoll fecal samples collected at inland sites (>100 km from the Atlantic coast) were categorized as “stopover” (Ohio, Pennsylvania, and Maryland, USA), whereas those collected along the US Atlantic coast (New Jersey, Massachusetts, and Rhode Island, USA) were categorized as “staging” (DeLuca et al., 2015; Morris et al., 2016; Warnock, 2010). Captured blackpolls were placed in a clean paper bag for 10 minutes to facilitate defecation and fecal samples were transferred to vials of Queen’s lysis buffer using sterile swabs. All fecal samples were kept frozen at −20ºC in the field for less than one month before long-term laboratory storage at −80ºC. Each sampled blackpoll was aged, sexed, weighed, and measured for subcutaneous fat deposits (scored 0-8) and wing chord. To assess energetic condition while controlling for structural size, as measured by wing chord, we calculated a Scaled Mass Index (SMI) following Peig & Green (2009).

### 16S rRNA amplicon profiling

DNA was extracted from 130 blackpoll fecal samples using the Qiagen PowerFecal DNA Kit (Qiagen, Hilden, Germany; 12830) before amplification and sequencing by the Genome Research Core of the University of Illinois at Chicago as previously described (Trevelline, MacLeod, et al., 2019). Briefly, polymerase chain reaction (PCR) was used to amplify a portion of the bacterial 16S rRNA gene for Illumina sequencing using the Earth Microbiome Project primers 515F (GTGCCAGCMGCCGCGGTAA) and 806R (GGACTACNVGGGTWTCTAAT) targeting the V4 region of microbial small subunit ribosomal RNA gene (Caporaso et al., 2011). Amplicon libraries were sequenced using a 2×251 paired-end run on an Illumina MiSeq. Additionally, we also sequenced 2 lysis buffer controls and 10 ‘blank’ extractions to control for microbial DNA found in commercial extraction kits (Salter et al., 2014). A total of 4,582,950 raw Illumina sequencing reads (mean of 32,274 per sample [n = 142] ± 3679 SE) were trimmed, paired, and quality filtered via the DADA2 pipeline (Callahan et al., 2016) in QIIME 2 (version 2021.11; Bolyen et al., 2019) using default parameters. Sequences that passed the quality filter were denoised into 1,774 amplicon sequence variants (ASVs), which were identified using the SILVA reference database (release 138; Quast et al., 2013). Identified ASVs were filtered to exclude sequences of non-bacterial origin (archaea, plants, eukaryotes, and mitochondria) before being passed to the R package *Decontam* (Davis et al., 2018) for the statistical identification and removal of 10 contaminant ASVs found in both control extractions and blackpoll fecal samples, reducing the total number of reads to 1,384,384 (mean of 10,987 per sample [n = 126] ± 1,200 SE) and 1,394 ASVs. We rarefied ASV tables to 273 sequences per sample (excluding 15 samples) before comparisons of alpha and beta diversity in QIIME 2. We used unrarefied ASV tables (excluding the 15 samples that failed to meet the rarefaction threshold) for analysis of taxonomic composition and downstream differential abundance testing (McMurdie & Holmes, 2014).

### Shotgun metagenomics

Extracted fecal DNA from breeding (Yukon Territory and British Columbia; n = 8) and staging blackpolls (Block Island; n = 12) were sequenced by CoreBiome, Inc. (St. Paul, MN) using shotgun metagenomics (BoosterShot™). Briefly, sequencing libraries were prepared using a procedure adapted from the Illumina Nextera Library Prep Kit (Illumina, 20018705) and sequenced on an Illumina NovaSeq using single-end 1×100 reads with the Illumina NovaSeq SP reagent kit (Illumina, 20027464). A total of 115,289,944 raw sequence reads (mean of 5,764,497 per sample (n = 20) ± 1,117,040 SE) were filtered for low quality (Q-Score < 20), trimmed of adapter sequences using cutadapt (version 2.10). To reduce the possibility of DNA contamination from hosts, reads were then mapped against the Yellow-rumped warbler genome (*Setophaga coronata coronata*; assembly mywa_2.1), the closest relative of the Blackpoll warbler with a fully sequenced genome, as well as the human genome (assembly GRCh38.p13) using bowtie2 (version 2.4.3). Reads that did not align to either genome were profiled for microbial functions using HUMAnN (version 3.0) and the UniRef50 database.

### Statistical analysis

Differences in metrics of microbiome alpha diversity (Faith’s phylogenetic diversity and ASV richness) were assessed using the non-parametric Kruskal-Wallis test with False Discovery Rate (FDR) correction in QIIME 2. Principal Coordinates Analysis (PCoA) using unweighted and weighted UniFrac distances (Lozupone & Knight, 2005) were used to test for differences microbiome community membership and structure across migration phases using permutational multivariate analysis of variance (PERMANOVA) and permutational analysis of dispersion (PERMDISP) in QIIME 2. Differences in the relative abundance of bacterial taxa across migration phases and metrics of avian body condition (SMI and fat scores) were tested in MaAsLin2 (analysis_method = “CPLM”) with age and sex as random effects and FDR corrections on variance-stabilized transformed data using arcsin(relative abundance^0.5^) (Mallick et al., 2021). Significant associations between migration phase, SMI, and fat score with HUMAnN 3.0 gene pathway abundances were identified using MaAsLin2 with negative binomial model (minimum prevalence = 0.5) and FDR corrections on cumulative sum scaled data. For all statistical analyses, P/Q-values ≤ 0.05 were defined as ‘significant’.

## RESULTS

### Blackpoll gut microbiome community composition and structure

We first investigated how blackpoll gut microbiota differed across three distinct phases of fall migration: breeding (n = 16), stopover (n = 55), and pre-migration staging (n = 40). For metrics of microbiome alpha diversity, there was a significant reduction in Faith’s Phylogenetic Diversity across phases of blackpoll migration (F = 14.30, P < 0.001; Fig. 1a). There were no differences in ASV richness across phases (F = 0.23, P = 0.890; Fig. 1b). For metrics of microbiome beta diversity, there were significant differences in both gut microbiome community structure (PERMANOVA pseudo-F = 7.35, P = 0.001; Fig. 2) and membership (PERMANOVA pseudo-F = 4.07, P = 0.001; Fig. S2) across phases of migration. Further, we also observed a progressive reduction of within-group microbiota dissimilarities (beta-dispersion) across migration phases for both microbiome community structure (PERMDISP F = 17.61, P = 0.001; Fig. 2c) and membership (PERMDISP; F = 7.68, P = 0.006; Fig. S2c), indicating an overall reduction of inter-individual microbiome variation as blackpolls migrate from breeding to staging habitats.

**Figure 1.**
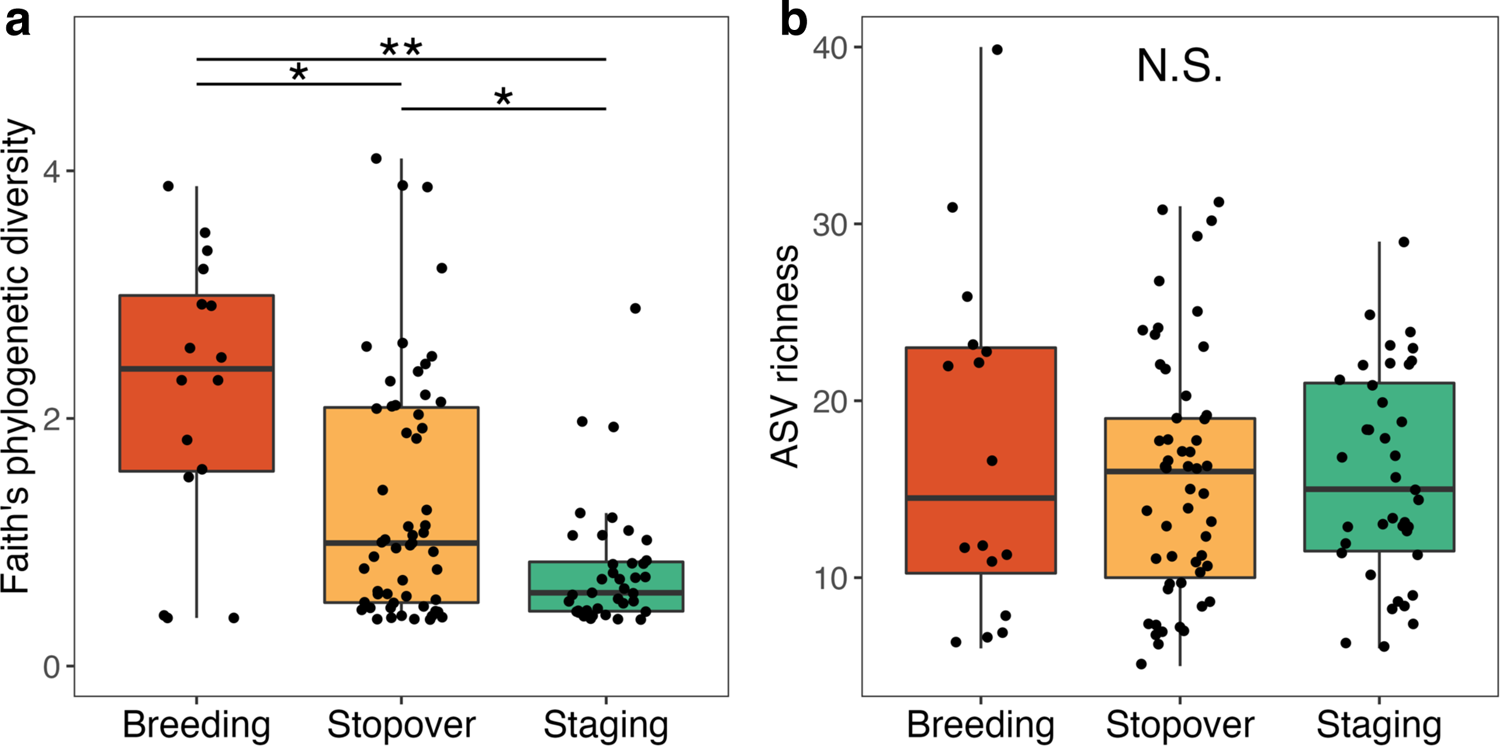
The Blackpoll warbler gut microbiome exhibited reduced alpha diversity over phases of fall migration. **(a)** Faith’s phylogenetic diversity differed significantly across phases of blackpoll migration. **(b)** Blackpolls exhibited no significant differences in ASV richness over phase of migration. ** denotes P ≤ 0.001 and * denotes P ≤ 0.05 after FDR correction. N.S. denotes no significant difference (P > 0.05).

**Figure 2.**
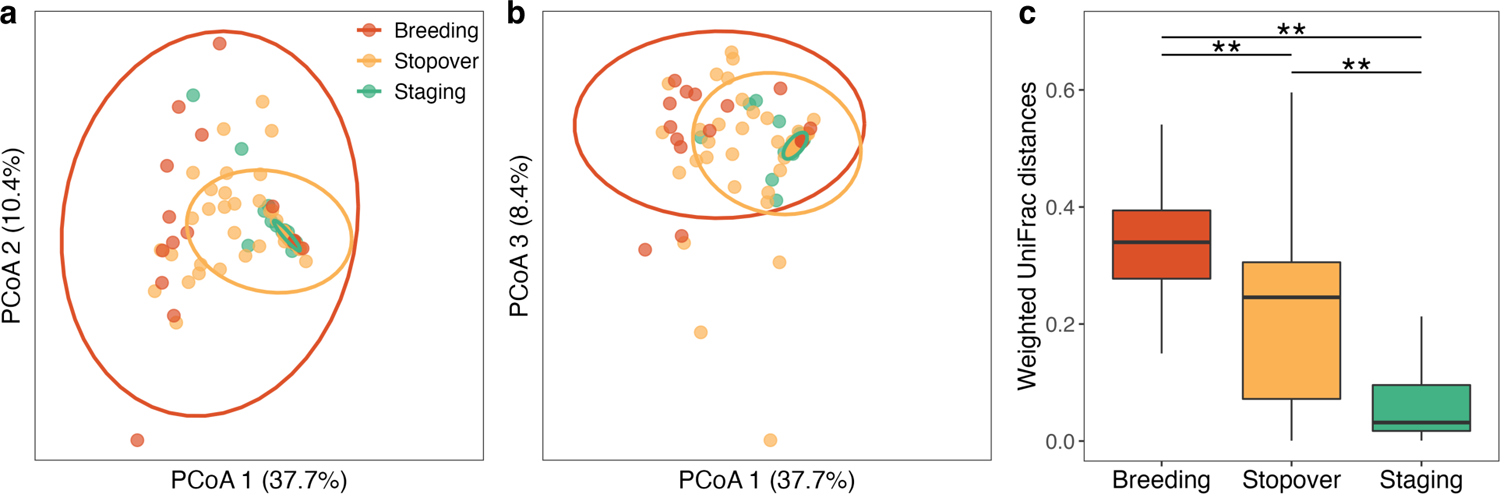
Convergence of the Blackpoll warbler gut microbiome community structure over phases of fall migration. **(a, b)** Principal Coordinates Analysis (PCoA) illustrating significant differences in microbiome community structure using weighted UniFrac distances (pseudo-F = 7.35, P = 0.001). **(c)** Within-group weighted UniFrac distances differed significantly across phases of blackpoll migration (PERMDISP F = 17.61, P = 0.001). ** denotes P ≤ 0.01 and * denotes P ≤ 0.05 after FDR correction.

Across all stages of migration, the blackpoll warbler microbiome was dominated by taxa in the phylum Proteobacteria (96%), followed by Firmicutes (3%) and 19 others with total relative abundances less than 1% (Fig. 3a; Supplemental Data S2). For phyla with relative abundances greater than 1%, Proteobacteria progressively increased as blackpolls migrated from breeding to coastal staging sites (Fig. 4a), while Firmicutes progressively decreased (Fig. 4b; Supplemental Data S2). At the family level, the blackpoll gut microbiome was dominated by three families of Proteobacteria: Enterobacteriaceae (54%), Yersiniaceae (23%), and Morganellaceae (9%; Fig. 3b; Supplemental Data S2). The most dominant bacterial family, Enterobacteriaceae (phylum Proteobacteria), progressively increased as blackpolls migrated to staging sites (Fig. 4c), and this pattern was almost entirely driven by a single ASV that could not be identified beyond the family level (Fig. 4d). There was no significant association between metrics of host body condition (SMI and fat score) with the relative abundance of bacterial phyla, families, or ASVs.

**Figure 3.**
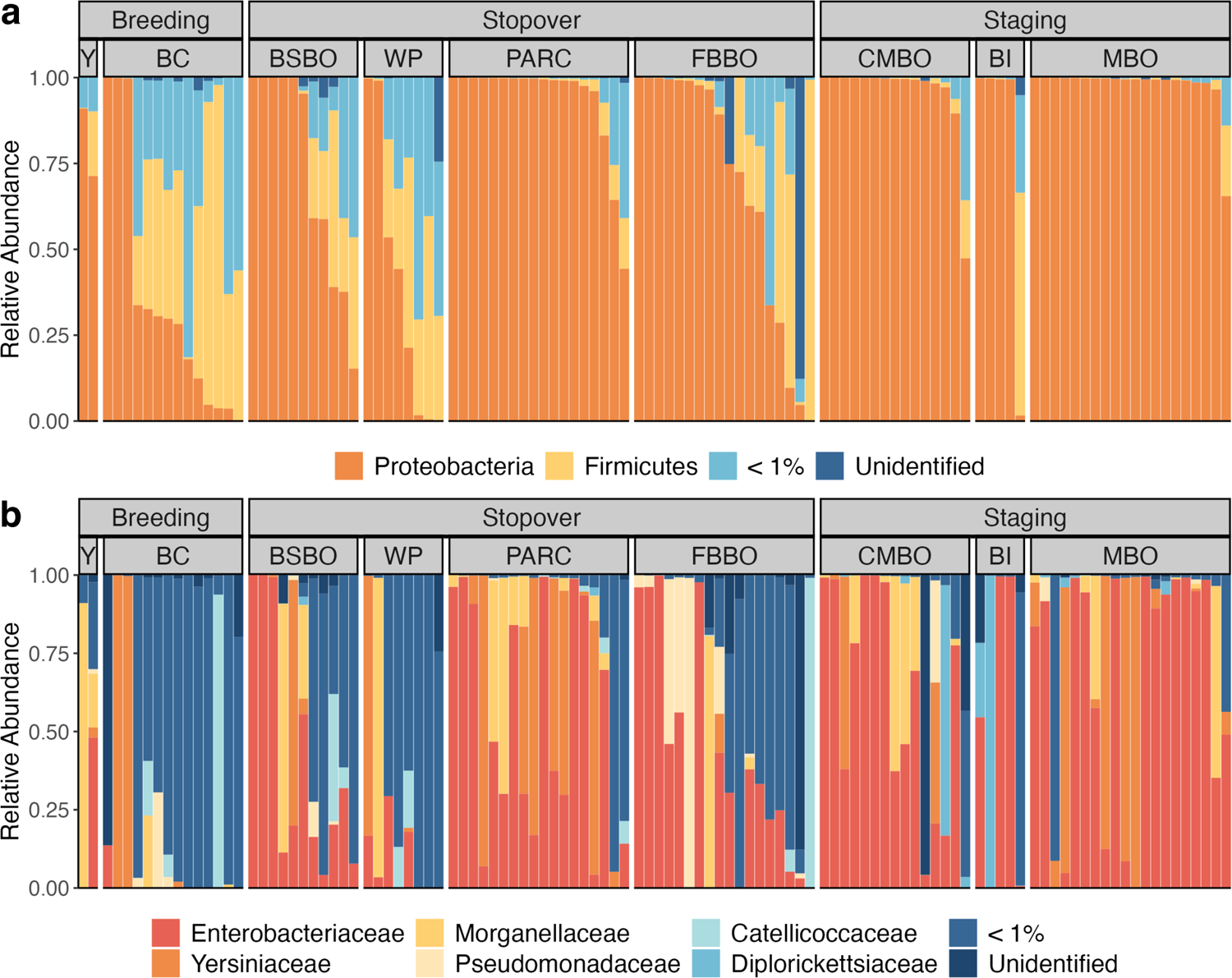
Remodeling of Blackpoll warbler gut microbiome composition over phases of fall migration. **(a)** Relative abundances of bacterial phyla and **(b)** families across phases of blackpoll migration. Letter codes denote sampling locations within stages of blackpoll migration. Breeding sites: Yukon Territory (Y) and British Colombia (BC); Stopover sites: Black Swamp Bird Observatory (BSBO), Winous Point (WP), Powdermill Avian Research Center (PARC), and Foreman’s Branch Bird Observatory (FBBO); Staging sites: Cape May Bird Observatory (CMBO), Block Island (BI), Manomet Bird Observatory (MBO).

**Figure 4.**
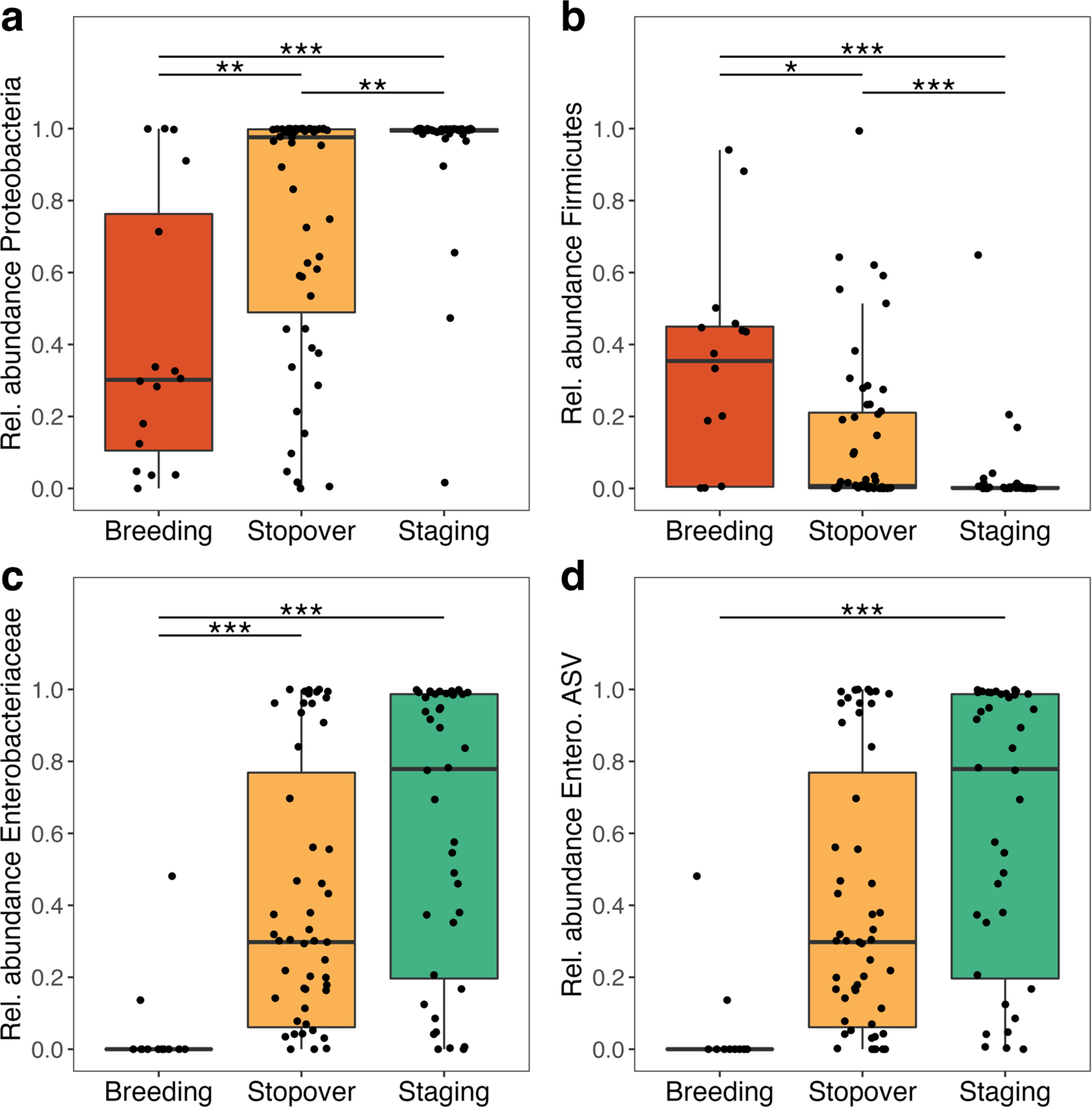
Taxonomic enrichment of Blackpoll warbler gut microbiota over phases of fall migration. The relative abundance of the bacterial phyla **(a)** Proteobacteria and **(b)** Firmicutes differed significantly over phases of blackpoll migration. The relative abundance of the bacterial family **(c)** Enterobacteriaceae and **(d)** a single ASV representing an undescribed species of Enterobacteriaceae differed significantly over phases of blackpoll migration. *** denotes P ≤ 0.001, ** denotes P ≤ 0.01, and * denotes. P ≤ 0.05 after FDR correction.

### Blackpoll gut microbiome function

Next, we used metagenomics to characterize differences in microbiome function between a subset of breeding (n=8) and staging (n=12) blackpolls. Our approach revealed a significant enrichment of 34 MetaCyc pathways between breeding and staging blackpolls (Fig. 5; Supplemental Data S3). The gut microbiome of staging blackpolls was enriched in a diversity of bacterial pathways including those involved in vitamin, amino acid, and fatty acid biosynthesis, as well as nucleoside/nucleotide cycling. Staging blackpolls were also enriched in carbohydrate degradation and homolactic fermentation of carbohydrates to the short-chain fatty acid lactate. In contrast, breeding blackpolls were primarily enriched in bacterial metabolic pathways involved in nucleotide degradation and polyprenoid biosynthesis.

**Figure 5.**
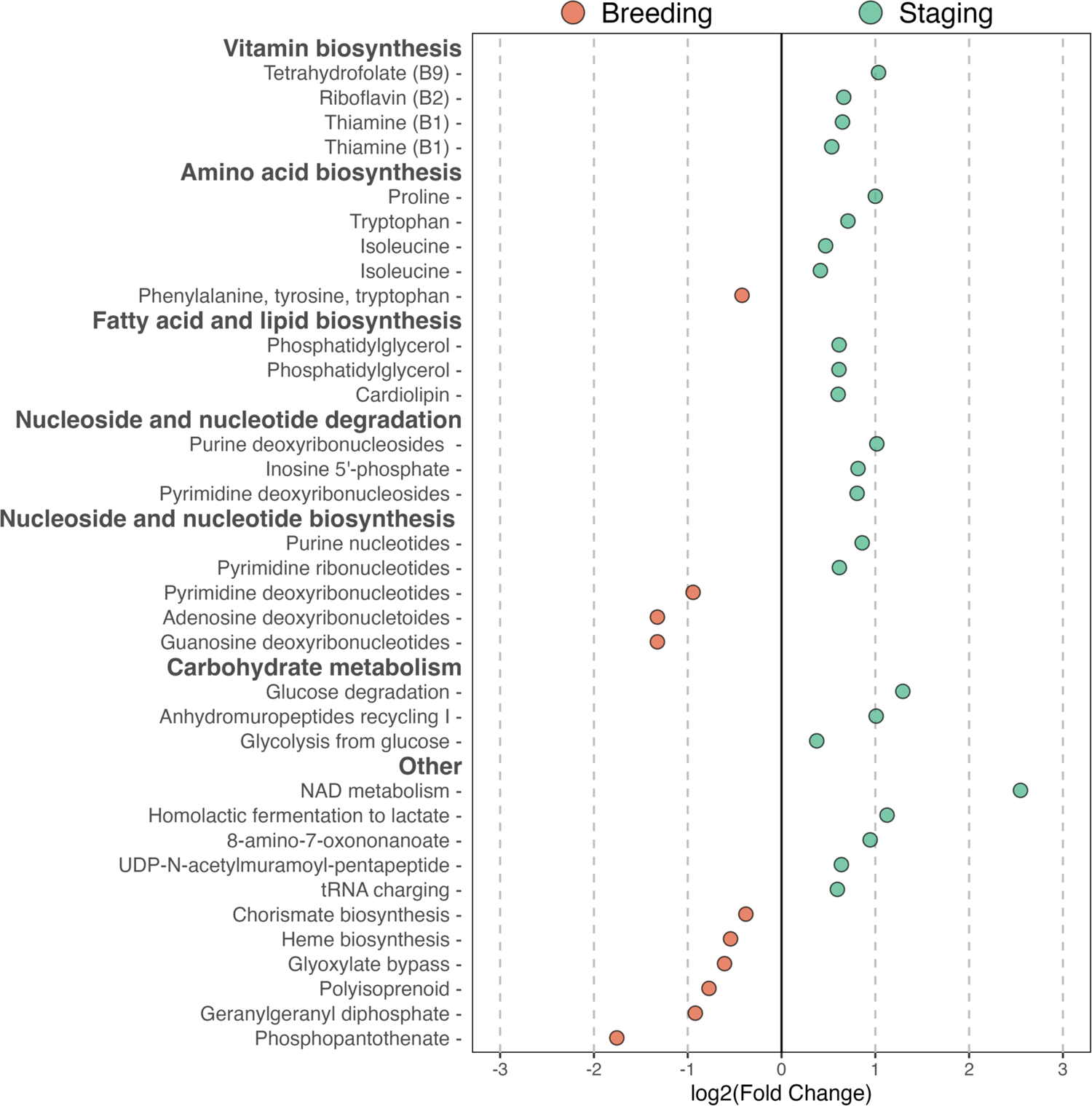
The gut microbiome of staging Blackpoll warblers is enriched in pathways related to energy metabolism. Log2 fold change plot illustrating 34 differentially enriched bacterial metabolic pathways between breeding (n = 8; red points) and staging (n = 12; green points) blackpolls. Red points represent pathways enriched in breeding blackpolls, while green points represent those enriched in staging individuals. All pathways were significant at the threshold of P ≤ 0.05 after FDR correction.

Lastly, we investigated whether bacterial pathways were associated with host body condition (SMI) and subcutaneous fat deposits. SMI and fat score were significantly associated with a total of 36 unique bacterial metabolic pathways (Fig. 6). SMI was positively associated with the enrichment of 16 pathways, primarily those involved in bacterial carbohydrate metabolism, as well as vitamin and amino acid biosynthesis (Fig. 6a). In contrast, fat scores were associated with the enrichment of 15 pathways, 3 of which were also positively associated with SMI: proline biosynthesis, sugar biosynthesis, and glycolysis (Fig. 6b).

**Figure 6.**
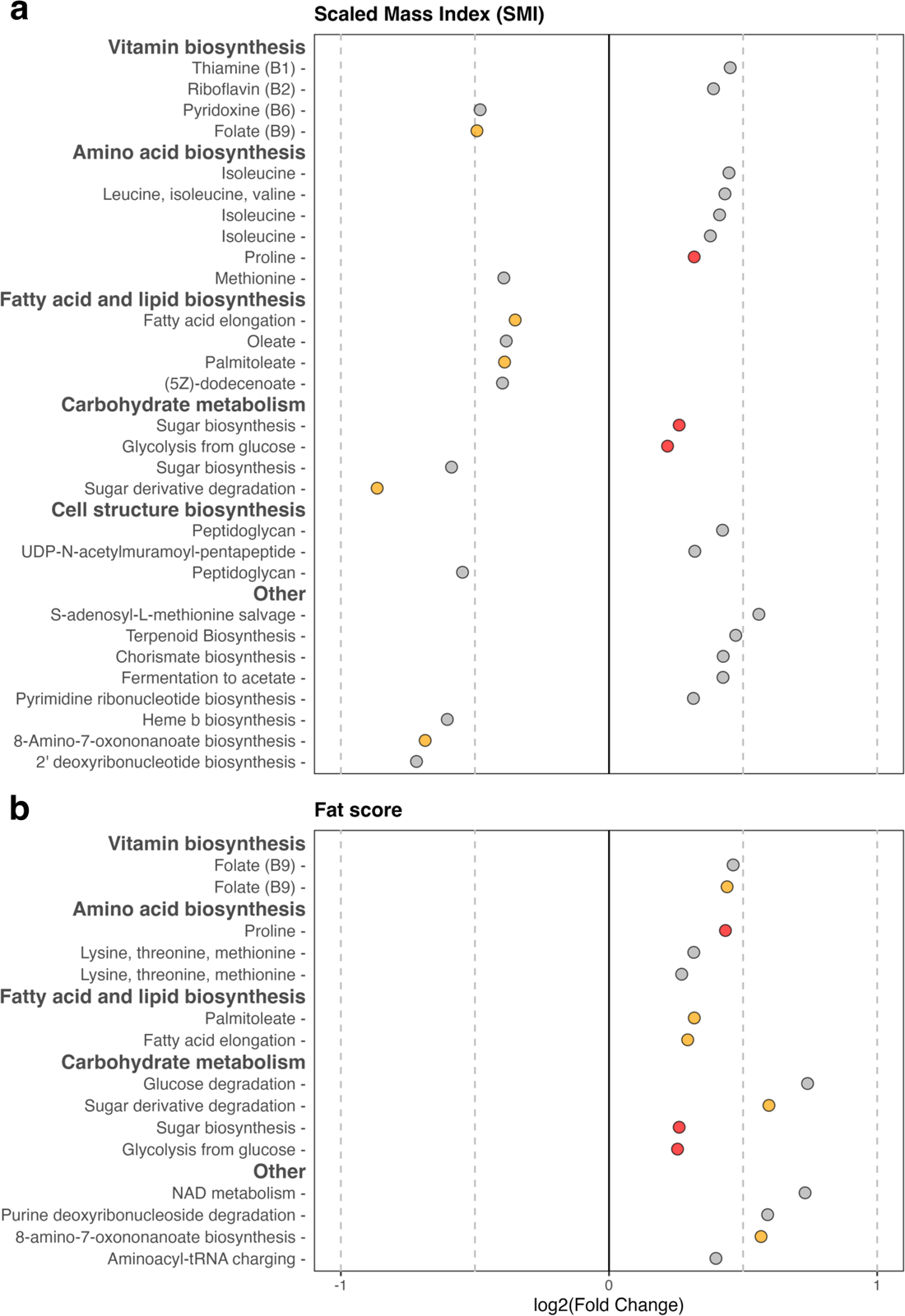
The enrichment of bacterial pathways is associated with metrics of blackpoll body condition. Log2 fold change plot illustrating 36 unique bacterial metabolic pathways between significantly associated with **(a)** Scaled Mass Index (SMI) and **(b)** subcutaneous fat scores. Yellow points represent bacterial pathways significantly associated with both SMI and fat, while red points represent those positively associated with both SMI and fat. All pathways were significant at the threshold of P ≤ 0.05 after FDR correction.

## DISCUSSION

In this study, we demonstrated that the Blackpoll warbler gut microbiome undergoes a dramatic and potentially adaptive remodeling during long-distance migration. Specifically, we showed that blackpoll gut microbiota exhibited a progressive reduction in phylogenetic diversity and within-group variability, as well as significant differences in composition across three distinct phases of migration. Notably, these patterns were driven by a significant increase in the relative abundance of Proteobacteria, and more specifically a single ASV belonging to the family Enterobacteriaceae. Metagenomic analysis revealed that the gut microbiome of staging blackpolls was enriched in several bacterial pathways with potential relevance for the metabolic needs of migrating blackpolls. Indeed, we found significant associations between several of these bacterial pathways and important indicators of energetic condition and migratory performance in blackpolls. Together, these results support the hypothesis that the gut microbiota of migratory birds is adaptively remodeled to meet the physiological and energetic demands of long-distance migration.

### Remodeling of microbiome community composition

We observed robust and significant response of the blackpoll warbler gut microbiota across phases of migration. While the dominance of Proteobacteria in the Blackpoll warbler gut microbiome is generally consistent with previous descriptions of wild passerine gut microbiota (Bodawatta et al., 2021; Grond et al., 2018), there were dramatic differences in microbiome community composition over phases of blackpoll migration. Consistent with previous studies showing that that migrating birds exhibit reduced gut microbial diversity (Risely et al., 2018; Skeen et al., 2021), blackpoll gut microbiota exhibited a progressive reduction in gut microbial phylogenetic diversity (but not ASV richness) from breeding sites in Western Canada to staging sites along the US Atlantic coast (Fig. 1). Moreover, blackpoll microbiome community membership (Fig. 2) and structure (Fig. S2) were distinct across migration phases, and were characterized by a progressive reduction in within-group variability across phases of migration. These findings suggest that the challenges imposed by long-distance migration may result in a convergence of microbiome composition mediated by natural selection favoring specific microbial lineages. Supporting this conclusion, differences in blackpoll gut microbial community composition were driven by dramatic shifts in the relative abundance of bacterial phyla belonging to Proteobacteria and Firmicutes among blackpolls at stopover and staging sites (Fig. 3). The increase in the relative abundance of Proteobacteria across phases of blackpoll migration was driven by a single ASV that belonged to the family Enterobacteriaceae (Fig. 3f), but whose species-level identification and function are unknown. The consistency across individuals with which this ASV came to dominate the microbiota in staging birds is consistent with the possibility that this ASV may harbor functions important for host migration performance. Notably, an increased relative abundance of Proteobacteria among migrating individuals was also observed in Kirtland’s warbler (*Setophaga kirtlandii*), a species closely related to the Blackpoll warbler (Skeen et al., 2021), suggesting that similar host factors (*e*.*g*., diet) during migration may result in convergent shifts in gut microbiota (Muegge et al., 2011). In contrast, our results differ from previous studies in *Calidris* shorebirds showing that migratory individuals were enriched in the bacterial family Corynebacteriaceae (phylum Actinobacteria; Risely et al., 2017, 2018), as the relative abundance of this family in blackpolls did not differ across migration phases. Cumulatively, these results suggest that microbiome remodeling may be a widespread phenomenon among migratory birds, but that the adaptive enrichment of potentially beneficial microbiota may be lineage specific.

### Enrichment of potentially adaptive microbiome functions

Compared to breeding individuals, the gut microbiome of staging blackpolls was enriched in 26 bacterial pathways, predominantly those involved in the bacterial biosynthesis of amino acids, vitamins, and lipids – all of which have been identified as important biomolecules for migrating birds (Jenni & Jenni-Eiermann, 1998; Klaassen, 1996). For example, staging migratory birds have been shown to accumulate fat-free mass in the form of muscle and certain organ tissues (Lindström & Piersma, 1993; Piersma, 1990), which can be catabolized (along with fat and glucose) as a source of energy during migration (Jenni & Jenni-Eiermann, 1998). Migrating birds primarily catabolize protein from pectoral muscle, digestive organs, and the gastrointestinal tract (Battley et al., 2000; Bauchinger & Biebach, 2001; Biebach, 1998; Jenni & Jenni-Eiermann, 1998; Schwilch et al., 2002b), leading to corresponding reductions in organ mass and function that must be recovered during stopover (Karasov & Pinshow, 2000; Muñoz-Garcia et al., 2012). The availability of plasma amino acids is the limiting factor in protein synthesis in birds (Scanes & Dridi, 2022) and previous work has shown that microbial amino acid synthesis can contribute to the host plasma amino acid pool (Metges, 2000). Alternatively, birds can also synthesize glucose from plasma amino acids via gluconeogenesis, providing an additional source of energy in fasting birds (Scanes & Dridi, 2022). In this study, the gut microbiome of staging blackpolls was enriched in the biosynthesis of several essential amino acids identified as critical for tissue growth in birds (Scanes & Dridi, 2022), and these pathways were also significantly associated with host body mass and subcutaneous fat deposits. While there are several other factors that mediate amino acid availability in migratory birds (e.g., Guglielmo, 2010), these results suggest the possibility that the microbiome of staging blackpolls provide a supplementary source of amino acids that can be used to rebuild catabolized digestive organs, which must be fully functional in order to digest and assimilate dietary nutrients during fall migration (McWilliams & Karasov, 2001). This hypothesis is supported by the positive association of several amino acid biosynthesis pathways and host body mass (Fig. 5).

The gut microbiome of staging blackpolls was also enriched in the biosynthesis of several B vitamins, most notably three separate pathways producing thiamine (Fig. 4). Thiamine (vitamin B1) is known to promote immune performance in vertebrates (Kumar & Axelrod, 1978), and thiamine deficiencies have been linked to disease susceptibility in wild birds (Balk et al., 2009). The energetic challenges of long-distance migration have been shown to reduce several metrics of immune performance in birds (Owen & Moore, 2006), raising the possibility that the enrichment of thiamine biosynthesis pathways among staging blackpolls is an adaptive property of the their microbiome that helps to maintain immune function under periods of intense metabolic demand.

Lastly, the gut microbiome of staging blackpolls was enriched in several bacterial pathways involved in cell membrane phospholipid biosynthesis. Phospholipids are readily digested and absorbed in the avian intestinal tract before being transported to the liver, where they are metabolized into triglycerides – the primary form of stored fat and the primary fuel source for long-distance migration in birds (Price, 2010). While it is tempting to speculate that microbial synthesized phospholipids could help facilitate fat storage in staging blackpolls, experimental evidence has demonstrated that dietary phospholipid content has no effect on body mass of migratory birds (Cerasale & Guglielmo, 2006). Regardless, the overall enrichment of microbial pathways with plausible contributions to migratory physiology among staging blackpolls supports the hypothesis that the remodeling of blackpoll gut microbiome confers adaptive advantages to hosts facing the energetic challenges of migration.

### Potential mechanisms

The remodeling of the gut microbiome among migrating blackpolls could be triggered by seasonal frugivory during migratory stopover. Many species of migratory birds (including blackpolls) shift from a protein-rich insectivorous diet during breeding to a carbohydrate-rich frugivorous diet during migration (Bairlein & Gwinner, 1994). The gut microbiome responds rapidly to changes in carbohydrate consumption (David et al., 2014; Smits et al., 2017), and recent work in bats provides evidence that fruit consumption can lead to the functional enrichment of carbohydrate degradation pathways (Ingala et al., 2021). In our study, staging blackpolls were enriched in bacterial metabolic pathways involved in sugar degradation, homolactic fermentation, and glycolysis, indicating that the gut microbiome of staging blackpolls may be specifically adapted to a fruit-based diet rich in dietary carbohydrates. Notably, the gut microbiome of staging blackpolls was enriched in the fermentation of fructose to the short-chain fatty acid lactate, which is readily absorbed in the avian intestine and essential to hepatic gluconeogenesis (Scanes & Dridi, 2022). Further, the association of host body condition with bacterial carbohydrate metabolism (sugar biosynthesis and glycolysis) suggest that these pathways may facilitate energy storage in migrating blackpolls (Fig. 5). While seasonal frugivory is thought to provide important source of carbohydrates for migratory birds, these fruits are comparatively low in protein (known as the “frugivory paradox”; Bairlein, 2002), which is necessary to rebuild catabolized digestive organs during stopover (Muñoz-Garcia et al., 2012). As noted above, the gut microbiome of staging blackpolls was enriched in several amino acids synthesis pathways, raising the possibility that gut microbiota may help meet the nutritional constraints of their diet during stopover and staging. Overall, these results suggest that the widespread consumption of carbohydrate-rich fruit among migratory birds could represent an additional adaptive response to the energetic demands of migration, but further study of how diet and microbiota covary across several migratory species is necessary.

Digestive systems are energetically-expensive to maintain, and previous work has established that phenotypic flexibility is an important mechanism by which migratory birds adaptively reduce certain digestive functions to help meet the energetic demands of long-distance flight (McWilliams & Karasov, 2001). Our results suggest that this paradigm can be extended to the gut microbiome, which we demonstrate exhibits remarkable flexibility in composition leading to several potentially adaptive functions during blackpoll migration. For instance, the enrichment of genes involved in bacterial protein and lipid metabolism over migration is consistent with the hypothesis that the gut microbiome acts as a phenotypically flexible trait that liberates the migratory birds from the energetic burden of maintaining these costly digestive processes. The proximate mechanisms driving the enrichment of these pathways could be initiated by changes in host diet and exposure to environmental microbiota, which are known to strongly shape the gut microbiome of migrants, and birds in general (Bodawatta et al., 2021; Grond et al., 2018). However, experimental techniques such as fecal microbiome transplants into gnotobiotic animal models (e.g., Sommer et al., 2016; Trevelline & Kohl, 2022) are necessary to definitively establish the functional effects of particular microbial taxa on host digestive physiology and migration performance.

### Conclusions

Overall, our study provides substantial evidence that the gut microbiome of a long-distance migratory bird undergoes a compositional and functional remodeling over fall migration. Whereas previous work on migrants has focused on compositional differences across just a few sites, we coordinated with researchers and avian monitoring stations to show that potentially important microbiome functions are enriched across several staging locations. These findings not only build upon numerous descriptive studies showing immense variation in gut microbiota among migratory birds (Bodawatta et al., 2021; Grond et al., 2018), but also contribute to a growing body of evidence that host-microbe interactions shape the ecology and evolution of wild vertebrates. Given that the energetic state of migratory birds is a critical determinant of migration performance, our results suggest that the gut microbiome of migratory birds may be an important yet unrecognized factor influencing the survival, fitness, and conservation of migratory birds.

## ACKNOWLEDGMENTS

We thank Lila Tauzer, Hilary Cooke, and Clara Cooper-Mullin for the collection of fecal samples for metagenomics sequencing at breeding and staging sites in Western Canada and Block Island, respectively. We also thank Trevor-Lloyd Evans of Manomet Bird Observatory, Annie Lindsay of Powdermill Avian Research Center, Mark Shieldcastle and staff of Black Swamp Bird Observatory, Liz Ames and staff of Winous Point Marsh Conservancy, Maren Gimpel of Foreman’s Branch Bird Observatory, David LaPuma of Cape May Bird Observatory for the collection of blackpoll fecal samples at stopover and staging sites. Blackpoll warbler range map provided by BirdLife International and Birds of the World. The authors have no conflicts of interest to declare.

## FIGURES

**Supplemental Figure S1.**
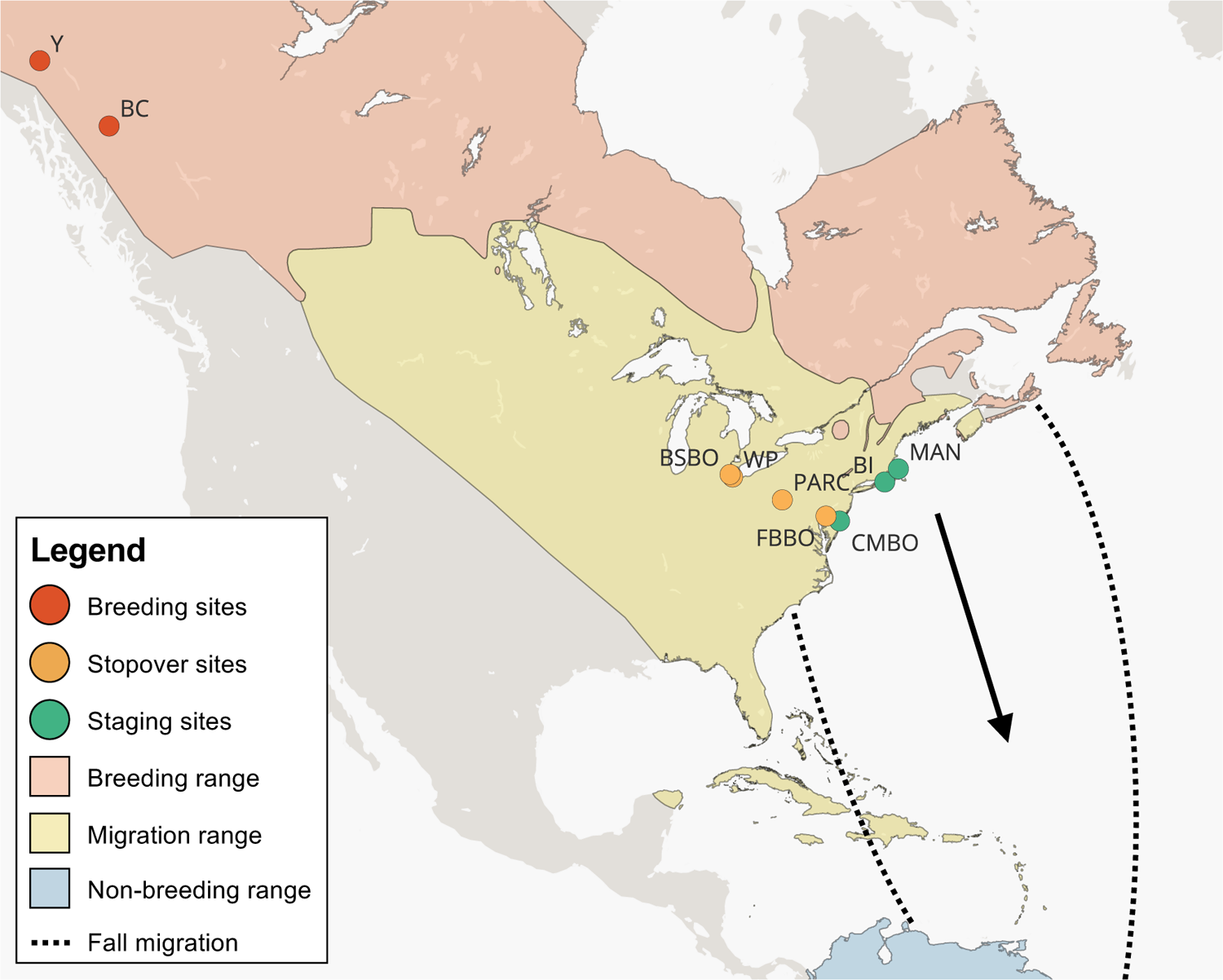
Sampling locations across phases of Blackpoll warbler fall migration. Points indicate sampling locations over three phases of blackpoll fall migration: Breeding sites (red points; n = 16) in Yukon Territory (Y) and British Colombia (BC); stopover sites (orange points; n = 55) at Black Swamp Bird Observatory (BSBO), Winous Point (WP), Powdermill Avian Research Center (PARC), and Foreman’s Branch Bird Observatory (FBBO); Staging sites (green points; n = 40) at Cape May Bird Observatory (CMBO), Block Island (BI), Manomet Bird Observatory (MBO). Blackpoll warbler range map provided by BirdLife International and Birds of the World.

**Supplemental Figure S2.**
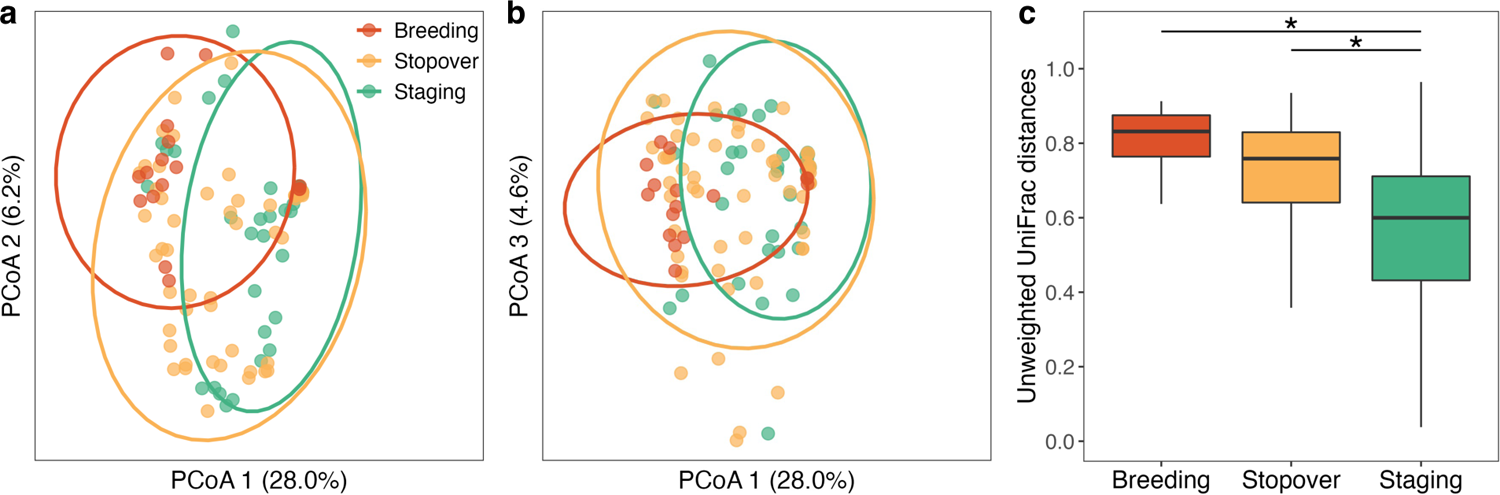
Convergence of the Blackpoll warbler gut microbiome community membership over phases of fall migration. **(a, b)** Principal Coordinates Analysis (PCoA) illustrating significant differences in microbiome community membership using unweighted UniFrac distances (pseudo-F = 4.07, P = 0.001). **(c)** Within-group unweighted UniFrac distances differed significantly across phases of blackpoll migration (PERMDISP; F = 7.68, P = 0.006). * denotes P ≤ 0.05 after FDR correction.

